# A phenotype-specific framework for identifying the eye abnormalities causative nonsynonymous-variants

**DOI:** 10.1101/2020.04.13.038059

**Authors:** Han-Kui Liu, Xiao Dang, Li-Ping Guan, Chang-Geng Tian, Sheng-Hai Zhang, Chen Ye, Laurent Christian Asker M. Tellier, Fang Chen, Huan-Ming Yang, Hao-Xiang Sun, Ji-Hong Wu, Jian-Guo Zhang

## Abstract

The most important role of variant pathogenicity predictors is to identify the disease-phenotype causative variant in studying monogenic diseases. In the last decade, machine-learning based predictors exhibited a relatively accurate performance for distinguishing the pathogenic variants and contributed a significant role for all disease-spectrums. Yet, few predictors can investigate the phenotypic significance of variants. Here we presented a phenotype-specific framework aimed to directly point out the phenotypic significance of predicted candidates, and showed its advancing performance in eye abnormalities. By training on eye-abnormalities causative variants, our method presented 96.2% accuracy, 96.1% precision, 93.4% recall for pathogenicity identification. Inconsistent with the modeling performance, identifying the single phenotype-causative variant from various sequencing variants is challenging for all predictors. Underlying the phenotype-oriented, our method significantly promoted the precision and reduced the cost for identifying the single causative variant from thousands of candidates. These advances highlight the significance of the phenotype-specific training method for studying disease.

## Introduction

Variant pathogenicity predictors based on machine learning model have been applied in studying disease comprehensively, such as CADD[1], FATHMM[2], MetaSVM[3], REVEL[4] and M-CAP[5]. Commonly, we apply these predictors to assist in the discovery of causative variants causing monogenic disease phenotypes[6]. A typical monogenic disease is caused by 1 or 2 rare variants among thousands of rare variants identified by genome sequencing[7], in which nonsynonymous variants contribute to the major component[8]. Limited annotation of nonsynonymous variants generally leads to a great challenge in assessing the pathogenicity, especially for most missense variants are termed uncertain significance (VUS) by ACMG guidelines[6]. These generic predictors gain knowledge from all disease-causal variants and provide powerful scores for estimating the variant pathogenicity. These scores displayed a relatively accurate classification in their training data which constituted by thousands of pathogenic variants and numeric-match controls. Inconsistent with their excellent modeling accuracy, identifying the single causative variant from hundreds of predicted candidates in practice is variable and may result in a severe precision. A poor precision in clinical test generates additional cost for verification, even leads to misdiagnosis. To tackle these plights, promoting the practical precision is necessary. Moreover, predictors that developed to cover all disease-spectrums provide basic and/or generic pathogenicity options, however, they remain unclear and/or nonspecific relationships between various candidate variants and observed phenotypes [9]. Few predictors provided options pointing to specific disease phenotype directly. Taken together with the practical precision and phenotypic significance, identifying the abnormal phenotype causative variant amongst a large number of candidates is challenging for all predictors. Recent studies showed advancing performance to investigate pathogenic variants using specific training data/features, such as disease-specific variants[10], variants in a gene[11] and gene family[12], tissue-specific gene expression[13], highlighting the importance for further development based on phenotype-specific data in this filed. Aimed to directly investigate the phenotype-causative variant and promote the practical precision, here we proposed a phenotype-specific framework and presented the performance and application in inherited eye disorders as an example.

## Materials and Methods

### Training set

To collect genes with eye abnormalities (EA) evidence, we obtained 2443 genes and the corresponding OMIM number related to eye abnormalities (HP:0000478) in the Human Phenotype Ontology (HPO) database. To select the pathogenic genes with clear molecular evidence, we focused on 696 OMIM numbers of inherited eye disorders from the Hereditary Ocular Disease (HOD) database, and an additional 49 OMIM numbers from our inhouse-knowledge mining. Using these 745 disease OMIM numbers (503 genes), we collected 11,956 pathogenic SNVs denoted as “Pathogenic” or “Likely pathogenic” in ClinVar, and “Disease” in UniProt/Swiss-Prot. These 11,956 pathogenic SNVs and corresponding 503 genes with phenotypic information of eye abnormalities and molecular information of inherited eye disorders were considered eye abnormalities causative.

For comparison, we selected 45,495 neutral SNVs presented in all 3 databases: 1000 Genomes Project, Wellderly[14], and VariSNP[15]. We annotated both causative and neutral SNVs with allele frequency and genetic consequence using VEP[16] and dbNSFP[17, 18] and remained 9,396 causative variants and 15,990 neutral variants met all following criteria finally: (1) minor allele frequency (MAF) < 1% in 1000 Genomes and gnomAD database; (2) nonsynonymous SNVs; (3) only present in one group of pathogenic SNVs or neutral SNVs. This final dataset was randomly divided into two subsets: four-fifths were used for training, while the remaining was used for testing.

### Phenotype specific training

Our method employed features from tissue-specific gene expression, gene-specific predictors and variant predictors. 30 human tissue expression were obtained from TiGER. 4 scores of gene predictors, RVIS [19], GDI[20], LoFtool[21], and SORVA[22], and 23 scores of 12 popular variant predictors, SIFT[23], PolyPhen-2[24], LRT[25], MutationTaster[26], MutationAssessor[27], FATHMM[28], PROVEAN[29], fitCons[30], PhyloP[31], GERP++[32], SiPhy[33] and phastCons[34] were obtained from the dbNSFP database. We selected the most serious one across all isoforms for any predictors with multiple protein isoforms in a given variant. In total, 80 features were assigned for training data. Missing features were imputed by the “Hmisc” R-package[35].

We trained the decision tree using “randomForest” R-package [36] and ensured the model accuracy followed a 5-fold cross-validation approach based on the training and testing dataset described above. In addition, we compared the performance of our method with other tools (REVEL, M-CAP, MetaSVM, VEST3[37], SIFT, CADD) on the testing dataset described above, and 2 validation datasets, novel gene set, control dataset described below. Accuracy was calculated by (TP+FP)/(TP+FP+TN+FN), precision was calculated by TP/(TP+FP) and recall was calculated by TP/(TP+FN), where TP refers to true positive, FP refers to false positive, TN refers to true negative and FN refers to false negative for each predictor. Performances of each predictor were assessed using the precision-recall curve.

### Validation sets

To evaluate the inherent bias of the training data, we employed two independent datasets: *validation set 1* and *validation set 2*. Variants presented in the training dataset or with MAF greater than 1% were excluded from these 2 datasets. *Validation set 1* included 928 rare inherited eye disorder pathogenic variants (labeled as pathogenic or likely pathogenic) and 6219 rare neutral variants (labeled as Benign or likely benign) newly reported in ClinVar after 06/02/2016. *Validation set 2* included 104 novel retina pathogenic variants reported in two recent studies [38, 39] and 1155 rare neutral variants in the UniFun database[40]. To ensure an accurate comparison for precision-recall curve analysis, we excluded variants with missing values in any predictors. After that, 186 pathogenic variants, 4010 neutral variants in *validation set 1*, and 104 retina pathogenic variants, 1155 neutral variants in *validation set 2* were remained.

### Novel gene set

To evaluate whether our method with the applicability to identify novel causative variants in novel genes, we used a novel gene set followed a 5-fold cross-validation approach. Training data was sorted by genes and divided into two parts: four-fifths of the genes and corresponding variants were used for training, while the remaining genes were considered as novel and used for validation.

### Control set

Our method specifically developed to predict eye abnormalities causative variants, and thus is expected to be limited in other disorders. To test the predictive ability upon other diseases, we collected the *BRCA1* and *BRCA2* variants as a control dataset. This dataset included 580 rare pathogenic, or likely pathogenic, germline nonsynonymous variants which cause familial breast cancer, and 229 rare benign nonsynonymous variants of *BRCA1* and *BRCA2*.

### Genome sequencing data

To evaluate the practical performance, we used the genome sequencing data of the Inherited Retinal Disease (IRD) project [38]. We accessed 767 CRAM/BAM files of 722 individuals from EGA (EGAD00001002656) with the approval of the author. We called variants using GATK [41] and annotated variants using VEP. We obtained 404 patients with clearly identified causative variants which were reported and verified in the study. From these 404 patients, we selected 204 patients and the corresponding 203 patients-phenotype causative nonsynonymous variants that absented in training data. These 203 causative variants were used for evaluating the practical precision and recall performance of our method and other predictors.

### Burden Analysis in genome-wide association study

Gene burden analysis was used to evaluate the association between the variant burden and phenotype. Gene carrying pathogenic mutations is considered a rare event, and the number of pathogenic mutations in one gene typically follows a Poisson distribution. Gene-based burden analysis was performed using the Poisson distribution test[42]. The expected pathogenic variants rate (M) was calculated as N/B, where N refers to all predicted damaging variants of all patients, B refers to the effective coding length of all genes. P-value was calculated as [1-Poisson cumulative distribution function (x-1, L×M)], where x refers to the observed number of predicted damaging variants of one gene, L refers to the effective coding length of the gene, and M refers to the expected pathogenic mutation rate. Gene with p-value accessed 0.05/2000 indicated a significant association with phenotype and suggested the heavy burden of variant contributing to the associated phenotype. We focused on the rare variants in the IRD population and remained 5992 variants met all the following criteria. (1) MAF < 0.05 in IRD population. (2) MAF < 0.005 in 1000 Genomes and gnomAD. (3) genotyping rate > 0.9. (4) nonsynonymous SNVs. (5) predicted damaging by our method. MAFs in population database and genetic consequences were annotated by VEP.

### Classification of causative genes

Logistic regression was used to compare the performance of features in the EA-causative genes classification, and to examine novel EA-related genes in whole genome-wide:

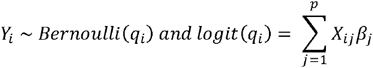

Where X refers to features of genes. Our feature was defined as: *Probability* (*G_i_*) = *D_i_*/(*D_i_* + *T_i_*), in which *D_i_* and *T_i_* refer to the predicted damaging (positive) numbers and predicted tolerance (negative) numbers in gene *i*. Y refers to gene-category, including EA-causative genes and tolerant genes. EA-causative genes refer to 503 genes used for model training. Tolerant genes refer to 1103 genes labeled as Non-Disease and Non-Essential from a previous study and with a frequency of loss-of-function (LoF) mutations greater than 5‰ either in 1000 Genomes or gnomAD databases. Other-genes refer to the remnant, which are neither EA-causative genes nor tolerant genes.

### Gene ontology annotation

We used gene ontology database for annotating EA -causative and -related genes. A hypergeometric distribution test was used for enrichment analysis. P-value was calculated as [1-Hypergeometric cumulative distribution function (x-1, m, n-m, k)], where x is the number of EA-genes in a cluster, m is the number of genes in a cluster in database, n is the number of genes in database, k is the number of EA-genes in database. A cluster refers to a group of genes with same GO-tag.

## Results

### Phenotype-specific training

To evaluate the performance of phenotype-specific framework for identifying causative variants, we focus on a well-studied category of diseases: the inherited eye diseases. Inherited eye diseases represent a vast spectrum of ocular genetic disorders generally developing eye abnormalities (EA) phenotype. Variants both with pathogenic information (causing inherited eye disorders in ClinVar) and phenotypic information (genes developed abnormal eye phenotypes in HPO and HOD) are considered as phenotype causative. Our phenotype-specific model for eye abnormalities (PSM-EA) was trained on 9,396 pathogenicity variants of 503 monogenic genes causing inherited eye diseases which developed abnormal eye phenotypes. (**Figure 1, Table S1**). A 5-fold cross-validation approach presented PSM-EA with an accuracy of 96.2% for pathogenicity identification. To demonstrate that our method is unbiased, we employed 2 independent validation sets (**Table S2**) and showed PSM-EA exhibited the best performance (**Figure 2**) compared with 6 widely employed predictors (REVEL, M-CAP, MetaSVM, VEST3, SIFT, CADD). The novel gene set presented PSM-EA is unbiased at gene-level and with the ability for discovering pathogenic variants in novel causative genes (**Figure 2**). The control set presented our phenotype-specific method (**Figure 2**) with a poor performance for eye abnormalities irrelevant diseases. These results indicated that PSM-EA is the most reliable and specific method for predicting pathogenic rare nonsynonymous variants that cause abnormal eye phenotypes. The accurate and phenotype-oriented classification provided confidence for pointing out the phenotypic significance of candidate variants.

**Figure 1:**
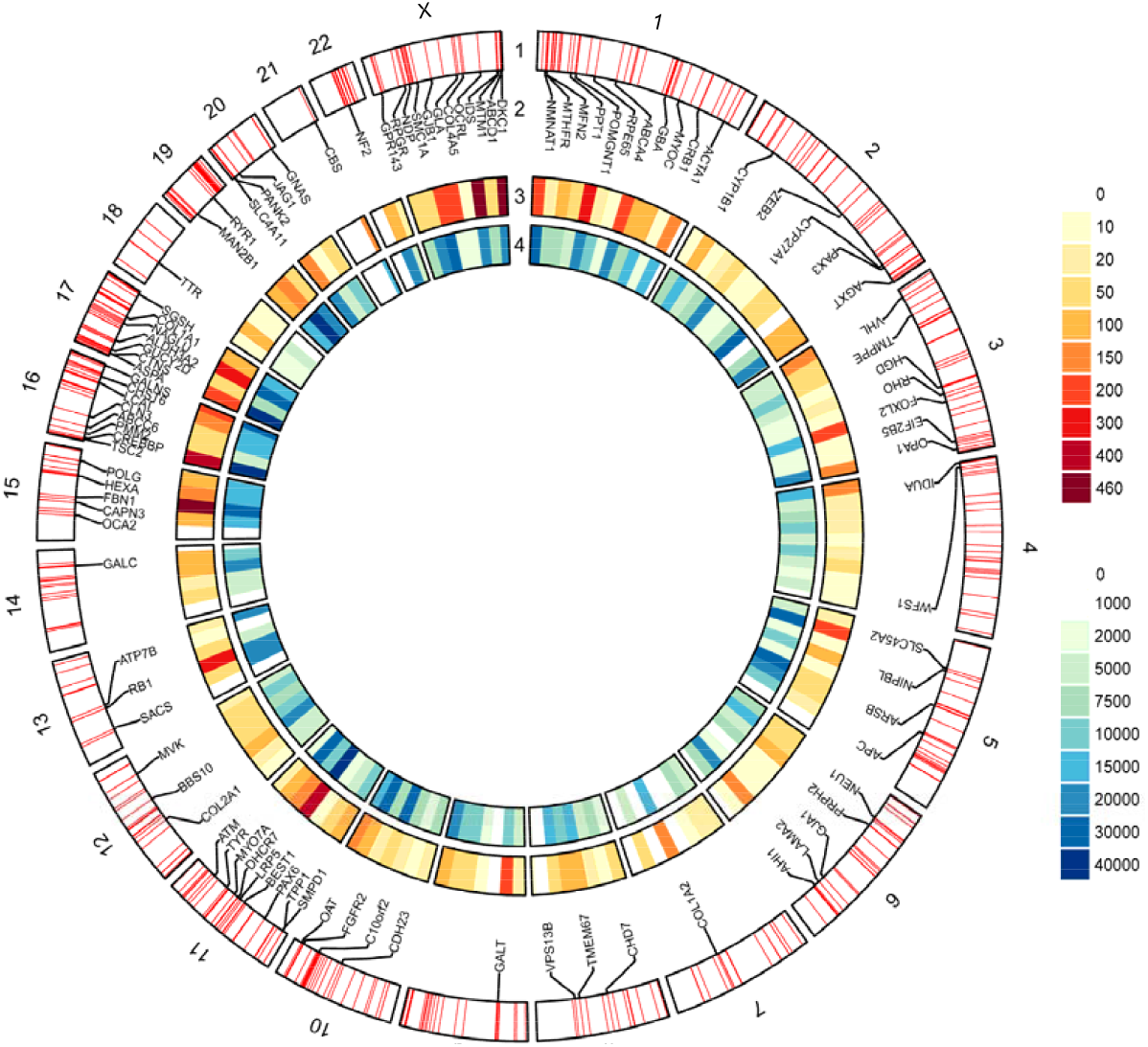
Summary of causative and predicted damaging nonsynonymous variants of eye abnormalities. (1) Ideograms of the 23 chromosomes of the human genome. The name of each chromosome is shown as label in the outermost circle. Eye abnormalities causative genes are shown as red lines. (2) The top 100 genes carry the most of causative nonsynonymous variants are shown as black lines and their corresponding labels. (3) The density of causative nonsynonymous variants is presented as the color within non-overlapping, 20-Mb windows. (4) The density of predicted damaging nonsynonymous variants is presented as the color within non-overlapping, 20-Mb windows.

**Figure 2:**
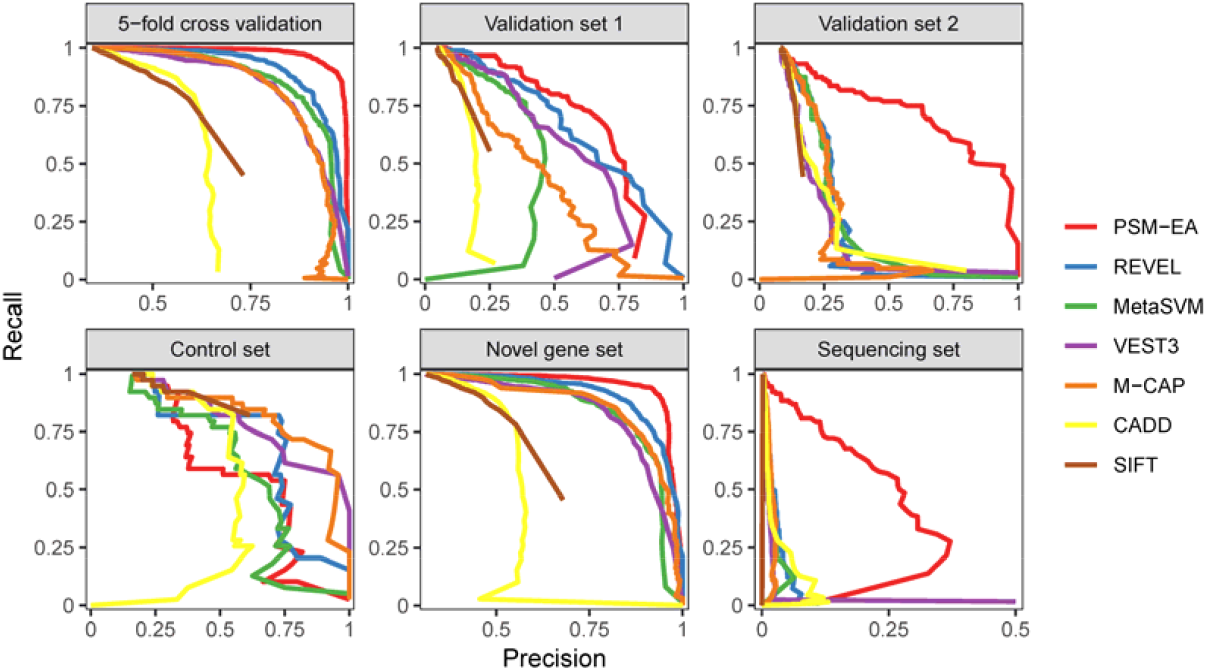
PSM-EA outperforms existing predictors. Precision-recall curves of PSM-EA, as well as comparison with other predictors in our testing dataset (5-fold cross-validation), independent validation datasets (validation set 1, validation set 2), non-ocular genetic diseases (control set), gene-based training (novel gene set) and genome sequencing data (sequencing set).

### Performance in genome sequencing practice

A patient with the monogenic disorder commonly carries the single causative variant, and the various predicted damaging variants generate further cost to identify the causative variant. Moreover, identifying the single causative variant among hundreds of candidates often result in severe practical precision even cause misdiagnosis. To estimate the practical precision in sequencing individuals, we used 203 verified causative variants of 204 IRD sequencing patients (**Table S3**). PSM-EA presented the enhanced performance (**Figure 2**), and the highest practical precision at the default setting (**Table 1**) and at the recall level of 0.5 (**Figure 3A**) for discovering causative variants. We noticed practical precisions are markedly different from modeling precisions in all tools. To our knowledge, precision is a significant measurement in modeling. To avoid the misunderstanding between modeling precision and practical precision, here we introduced the cost measurement for estimating the cost for identifying a pathogenic variant from sequencing patient and computed the cost as N_PD_/N_C_, in which N_PD_ refer to the number of predicted damaging variants in all 204 patients and N_C_ refer to the number of predicted damaging causative variants. Using the default setting of predictors, we showed PSM-EA with the lowest cost (**Figure 3B**) for identifying the single causative variant of sequencing patient. Generally, PSM-EA restricted an average of 15 predicted damaging variants in sequenced individuals and significantly (P-value of t-test = 7e-129) narrowed down 72-95% of unrelated variants in comparison with other predictors (**Figure 3C**). These results indicated that the phenotype-specific method promoted the precision-recall performance and reduced the cost for practical pathogenicity identification significantly.

**Table 1.** Practical performance on 204 sequencing IRD-patients with 203 novel causative variants

| Predictors | Recommended threshold | Modeling precision<sup>a</sup> | Prediction of causative variants |  |  | Predicted damaging variants |  |  | Practical performance |  |  |
| --- | --- | --- | --- | --- | --- | --- | --- | --- | --- | --- | --- |
|  |  |  | Damaging | Tolerated | Unknown | Total<sup>b</sup> | Median<sup>c</sup> |  | Recall<sup>d</sup> | Precision<sup>d</sup> | Cost<sup>e</sup> |
| PSM-EA | >0.5 | 0.961 | 116 | 15 | 73 | 2,634 | 15 |  | 0.885 | 0.044 | 22.707 |
| REVEL | >0.5 | 0.823 | 91 | 32 | 80 | 8,273 | 54 |  | 0.740 | 0.011 | 90.912 |
| MetaSVM | >0 | 0.821 | 83 | 40 | 80 | 8,399 | 58 |  | 0.675 | 0.010 | 101.193 |
| VEST3 | >0.5 | 0.624 | 100 | 23 | 80 | 21,242 | 175 |  | 0.813 | 0.005 | 212.420 |
| M-CAP | >0.025 | 0.596 | 115 | 3 | 85 | 26,191 | 191 |  | 0.975 | 0.004 | 227.748 |
| CADD | >20 | 0.523 | 188 | 15 | 0 | 39,086 | 301 |  | 0.926 | 0.005 | 207.904 |
| SIFT | <0.05 | 0.491 | 104 | 17 | 82 | 34,977 | 282 |  | 0.860 | 0.003 | 336.317 |

<sup>a</sup> Modeling precision are computed using our testing dataset.
<sup>b</sup> All unique predicted damaging variants in 204 IRD-patients
<sup>c</sup> The median of predicted damaging variants in 204 IRD-patients.
<sup>d</sup> Recall and Precision are computed as $TP/(TP+FN)$ and $TP/(TP+FP)$ , where TP refers to the number of predicted damaging in causative variants, FN refers to the number of predicted tolerant in causative variants, FP refers to the number of predicted damaging in non-causative variants.
<sup>e</sup> Defined as the cost for identifying the single causative variant in sequencing patient, and computed as $N_{PD}/N_C$ , in which $N_{PD}$ refers to the number of predicted damaging variants in all 204 patients and $N_C$ refers to the number of predicted damaging causative-variants.

**Figure 3:**
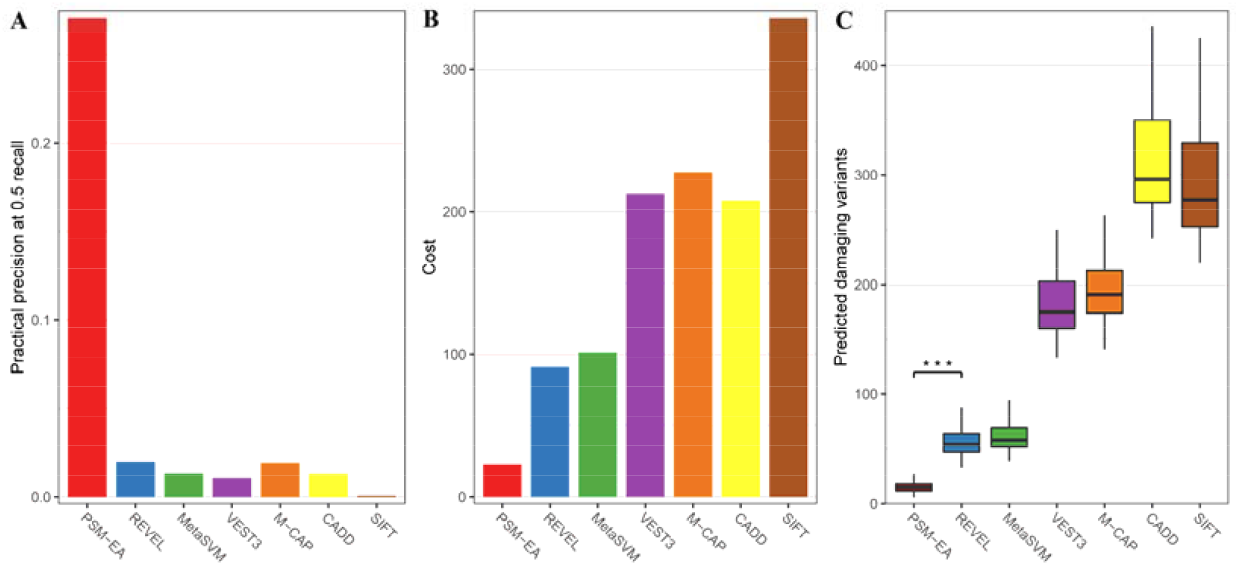
Practical performance in 204 sequencing IRD-patients. A total of 203 novel causative variants of 204 sequencing IRD-patients were used to evaluate the practical performance. (A) The precisions at recall of 0.5 present PSM-EA with the best performance. The Cost (B) was defined as the cost for identifying the single causative variant in sequencing patient, and computed as N_PD_/N_C_, in which N_PD_ (C) refers to the number of predicted damaging variants in all 204 patients at default setting and N_C_ refers to the number of predicted damaging causative-variants.

We also observed a similar performance for identifying 101 causative variants of a Chinese retinitis pigmentosa study [43, 44]. These replications suggested PSM-EA is robust across different ancestral populations (**Table S4**).

### Application in the association study

To extend the application in studying disease, we applied PSM-EA with gene burden analysis for evaluating the burden of predicted damaging rare variants of each gene in the IRD dataset. Statistical evidence showed 25 genes harbored predicted-damaging variants more than expectation significantly (**Table S5**), in which 24 genes existed in 503 causative genes and the novel gene *RBP3* was reported with critical function for vision in a recent study[45]. The lambda genomic factor 0.45 indicated no inflation of statistic method and/or no stratification of population. These results demonstrated PSM-EA performing well with the statistical method for identifying high confidence causative genes.

### A comprehensive reference of eye abnormalities causative variants and genes

A key goal of genetic research is to identify disease causative variants/genes and to study its functional roles in causative-phenotypes. Previous methods, such as family-based and population-based, were limited in the pedigree information or statistical power and resulted in sporadic candidates. To provide a comprehensive annotation of inherited eye disorders, we generated a map of 4,378,769 unique predicted damaging variants from all rare nonsynonymous variants in the human genome using PSM-EA (**Figure 1**). In addition, to provide a supplement for the spectrum of EA-causative genes, we employed a logistic regression based on a newly defined feature, the proportion of predicted damaging variants per gene, for evaluating the probability of EA-causative gene in genome-wide. This feature showed significant differences between EA-causative genes and tolerant genes (**Figure 4A**), and with an accurate of 90.3% better than the other 3 gene-features for classification of causative genes (**Figure 4B**). Using this feature, we examined 1527 novel EA-related genes (**Table S6**) from 19238 other genes with a false positive rate of 1.63%. To explore the function of these EA-related genes, we performed the gene ontology (GO) annotation and HPO annotation. GO annotation exhibited a group of genes playing a role (**Figure 5**) in eye development. HPO annotation exhibited 506 related genes with eye abnormalities.

**Figure 4:**
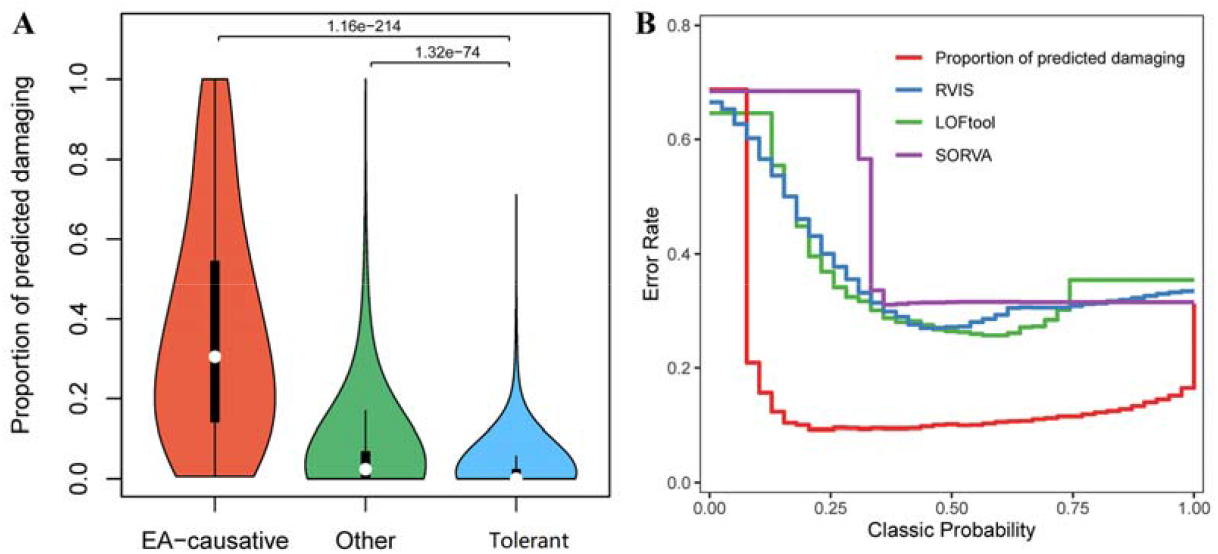
New feature for classifying eye abnormalities causative genes. The significant difference of distribution of our newly defined feature (Proportion of predicted damaging variants) among 3 gene groups: EA-causative genes, Tolerant genes and Other genes (A) and the comparison of error rate of classifying EA-causative genes using our feature (red) and other 3 gene features (B).

**Figure 5:**
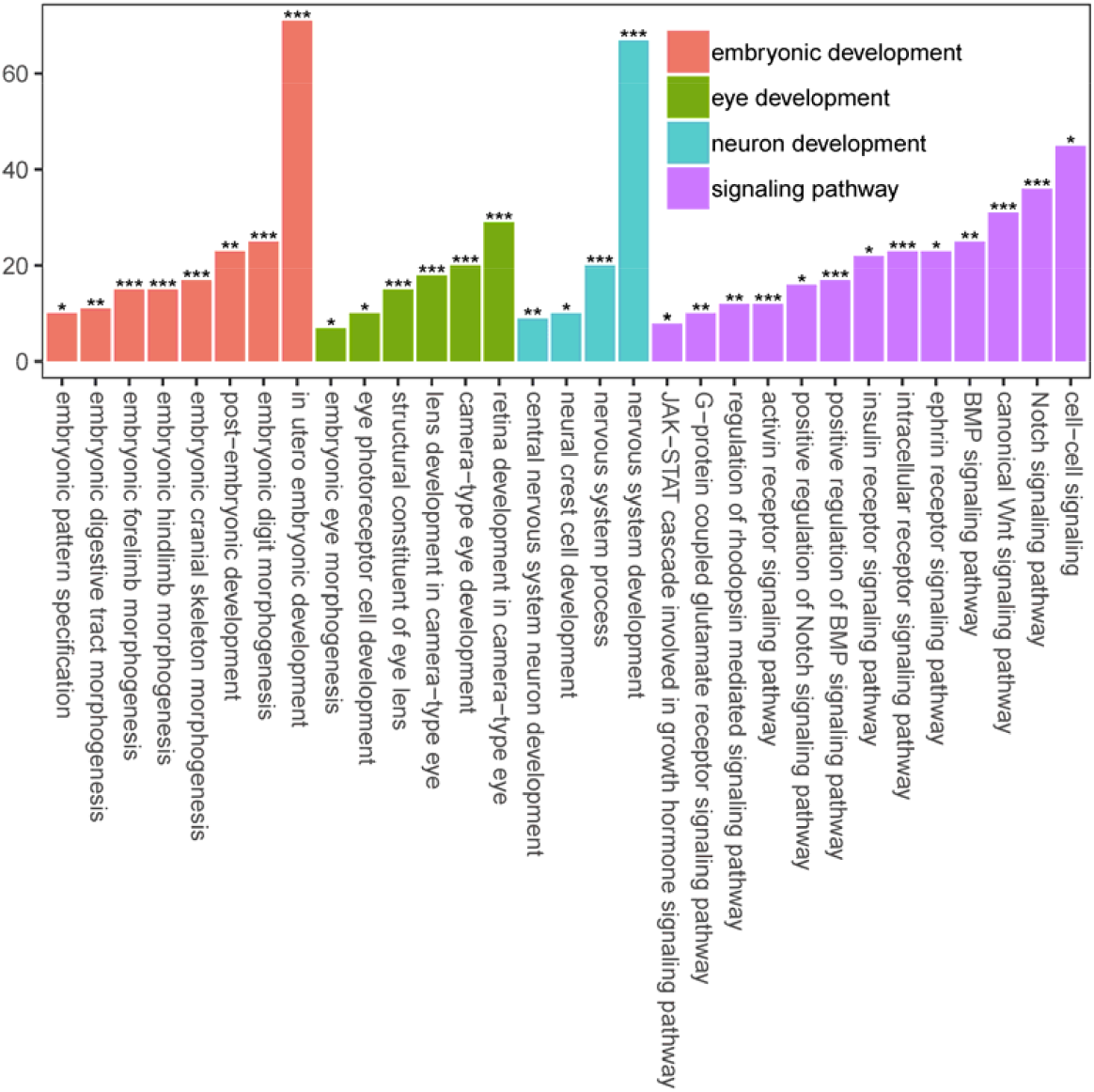
Gene ontology analysis. Bar refers to the number of genes in a cluster. Significance refers to the p-value calculated by hypergeometric distribution test and adjusted by FDR method, in which * refers to p-value <0.05, ** refers to p-value <0.01, *** refers to p-value <0.001. these results present the EA-related genes play a role in embryonic, eye, neural development and signaling pathway.

## Discussion

Supervised machine-learning based method was widely applied in identifying disease causative variants. Importantly, the characteristic of training data determined the performance and significance of a model. Predictors developed for all diseases causative variants represent generic pathogenicity. A recent study showed the performance of predictors was highly variable and differed by disease phenotypes[46]. We believe the phenotype-variable performances were determined by an underlying cause, that is the characteristics are differed by disease phenotypes, and the heterogeneous of training characteristics lead to variable predictions. We performed an unsupervised principal component analysis on 13 different diseases with the most pathogenic variants in ClinVar and their corresponding 80 features. We used the first 2 principal components for representing the phenotype-characteristics and for estimating the characteristics-heterogeneity. ANOVA showed significantly different diseases-characteristics among groups (**Figure S1**), which suggests a replicate conclusion that training a model based on the disease- and/or phenotype-specific data is necessary and advisable[10–13, 46]. For this purpose, we presented a phenotype-specific framework, building a machine-learning classifier based on known pathogenic variants of a specific disease phenotype. On this basis, we developed a random forest classifier for estimating the pathogenicity of rare nonsynonymous variants causing abnormal eye phenotypes. Importantly, we demonstrated that phenotype-specific classifiers significantly promote the practical precision, reduce the cost for the identification of causative variants and directly point out the causing phenotype for candidates. Moreover, we provided a comprehensive annotation for eye abnormalities related variants/genes. With the service of our method, researchers can obtain directive annotation and focus on the most significant variants among various VUS labeled by ACMG guidelines. These advances are critical for promoting the diagnosis rate in the clinical test and assisting in the discovery of causative variants for a specific phenotype. With the advancing performances, phenotype- and disease-specific framework showed great research prospects and may lead to the development of pathogenicity classifiers in the next period.

## RESOURCES TABLE

| Database/tools | Source |
| --- | --- |
| HPO | https://hpo.jax.org/app/download/annotation |
| Hereditary Ocular Disease | http://disorders.eyes.arizona.edu/physicians/clinical-descriptions |
| ClinVar | https://www.ncbi.nlm.nih.gov/clinvar/ |
| OMIM | https://omim.org/ |
| UniProt/Swiss-Prot | https://www.uniprot.org/ |
| TiGER | http://bioinfo.wilmer.jhu.edu/tiger/ |
| 1000 Genomes project | https://www.internationalgenome.org/ |
| gnomAD | https://gnomad.broadinstitute.org/ |
| GO | http://geneontology.org/ |
| PSM-EA | https://github.com/danria/PSM-EA |

## Acknowledgements

This study was supported by Shenzhen Municipal of Government of China (JCYJ20170412153248372 and JCYJ20180507183615145), The National Key R&D Program of China (NO.2016YFC1305900 and NO.2017YFC1308402).

We thank the NIHR BioResource, University of Cambridge, and the NIHR BioResource Rare Diseases BRIDGE consortium, which has sequenced and shared the inherited retinal disease data used in this analysis.

## Author contributions

H.L. and X.D. contributed equally. J.Z. and H.L. designed the study. X.D., L.G., S.Z., C.Y. and J.W. collected the genetic data. H.L., X.D., C.T and H.S. performed bioinformatic and statistical analysis. J.Z., L.G. and L.C.A.T. revised the manuscript. J.Z., J.W., H.Y. and F.C. supervised the research. All authors had full access to the final version of the manuscript and agreed to the submission.

---

**Figure S1:**
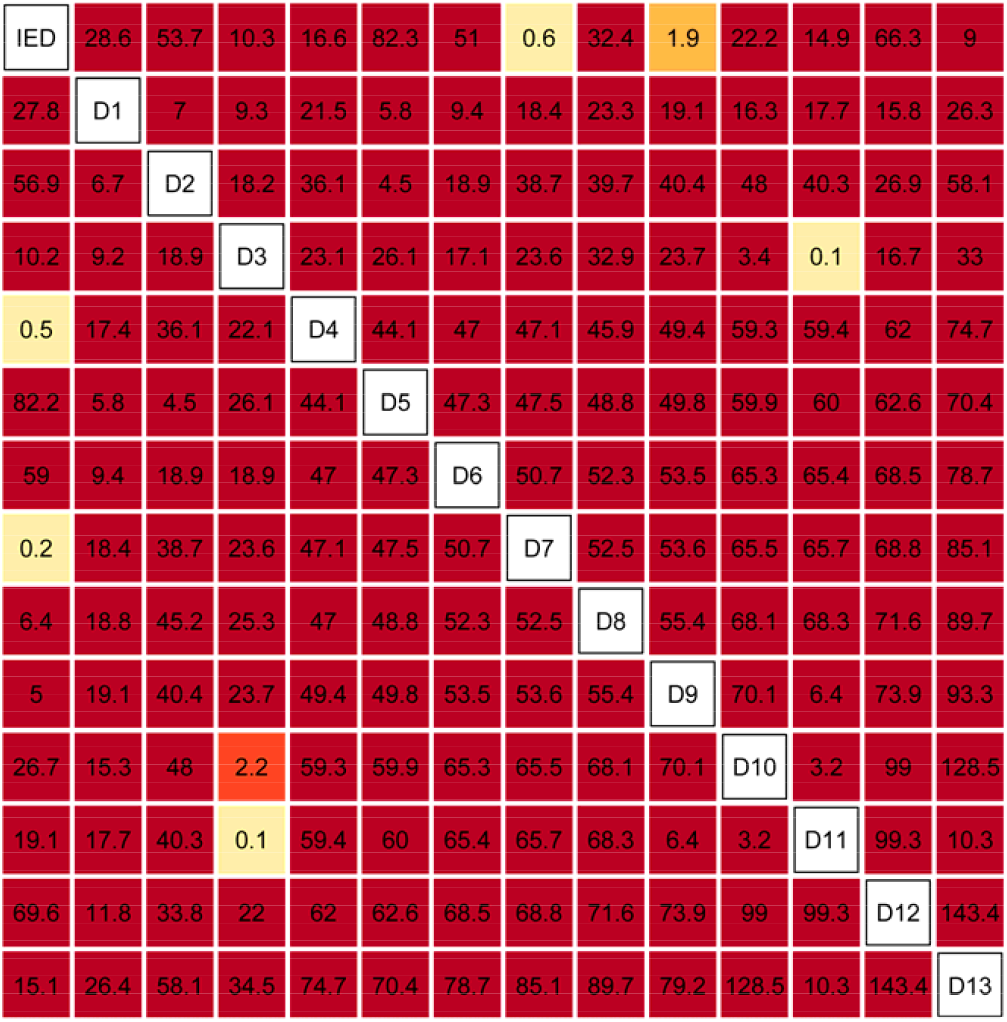
Diseases differ characteristic. Kruskal-wallis one-way ANOVA present significantly differenct components of 14 diseases characteristic. Components were calculated from the 80 features of disease variants using principal component analysis. IED refers to Inherited eye disorders. D1-D13 refer to other 13 diseases with the most pathogenic variants in ClinVar (1, Cardiovascular phenotypes; 2, Familial hypercholesterolemia; 3, Severe myoclonic epilepsy in infancy; 4, Primary dilated cardiomyopathy; 5, Ciliary dyskinesia; 6, Lynch syndrome; 7, Ehlers-Danlos syndrome, type 4; 8, Primary pulmonary hypertension; 9, Cystic fibrosis; 10, Marfan syndrome; 11, Pseudoxanthoma elasticum; 12, Duchenne muscular dystrophy; 13, Hereditary factor VIII deficiency disease). Numbers in the tile refer to –log10(p-value). Colors in tile refer to significance after FDR adjustment (light yellow, un-significant; yellow, p-value < 0.05; red, p-value < 0.01; Dark red, p -value < 0.001). Upper triangle: the first principal component. Lower triangle: the second principal component.

## Notes

### Competing Interest Statement

The authors have declared no competing interest.

